# Echocardiographic mitral valve association with morphometric measurements in Cavalier King Charles Spaniels via Inverse Probability Weighting analysis

**DOI:** 10.1101/2021.04.15.439951

**Authors:** Mara Bagardi, Sara Ghilardi, Chiara Locatelli, Arianna Bionda, Michele Polli, Claudio M. Bussadori, Fabio M. Colombo, Laura Pazzagli, Paola G. Brambilla

## Abstract

Development and progression of myxomatous mitral valve disease (MMVD) in Cavalier King Charles Spaniels (CKCS) are difficult to predict. Identification at a young age of dogs with a morphotype associated with more severe mitral disease is desirable. The aims of this study were to: 1) describe the physical, morphometric, and echocardiographic features of MMVD affected Cavalier King Charles Spaniels (CKCS) in American College of Veterinary Internal Medicine (ACVIM) class B1; 2) evaluate the influence of morphometric physical measurements on murmur intensity, mitral valve prolapse (MVP), regurgitant jet size and indexed mitral valve and annulus measurements. Fifty-two MMVD affected CKCS in ACVIM class B1 were included. This is a prospective clinical cross-sectional study. Morphometric measurements, which included body, thorax, and the head sizing of each dog, have been investigated to establish the association with heart murmur intensity, valvular and annular echocardiographic measurements, MVP and regurgitant jet size using inverse probability weighting (IPW) analyses to adjust for confounding. The IPW analyses showed that when head length and nose length decreased, dogs had more severe regurgitant jet size. Furthermore, subjects with more pronounced head stop angle had thicker anterior mitral valve leaflets. A brachycephalic morphotype, with dogs more similar to King Charles Spaniel breed in cephalic morphology, is associated with a more severe regurgitant jet size and with valvular characteristics related to worse forms of MMVD.

## Introduction

Myxomatous mitral valve disease (MMVD) is the most common acquired cardiac disease in canine patients [1]. Some studies indicate a polygenically inherited component of the disease, and at least in some of the highly susceptible breeds, early predictors of MMVD development and progression, such as morphotype, could allow for improved breeding recommendations [2-6]. As reported by Parker et al., the same genes can affect both dogs’ body size and heart development [7]. In fact, it is demonstrated that small breed dogs are more predisposed to MMVD development, especially Cavalier King Charles Spaniels (CKCS) [8]. Pedersen et al. (1999) showed that there is a negative correlation between body weight and mitral valve prolapse (MVP) in this breed [9]. Furthermore, in the CKCS, MMVD is associated with earlier onset and thus with potentially greater cardiac morbidity and mortality compared to other breeds [8,10,11]. Yet, the preclinical period often varies markedly among subjects, making it challenging for clinicians to identify those that will eventually develop clinical signs [12,13]. For these reasons, the early identification of a morphotype associated with a more severe MMVD can have several advantages. This could allow clinicians to monitor the dogs in a very targeted way and to educate breeders in the selection of subjects without some phenotypical characteristics related to a more severe MMVD and/or more rapid progression of the disease. This could be possible in the context of a breeding selection program, which should also consider all the heritable disorders of the CKCS.

To the best of our knowledge, only one study evaluated the prevalence and severity of MVP related to the size of the thorax in dogs, and in particular in Dachshunds,3 but no study has ever analyzed the association between MVP or MMVD severity and morphometric measurements in CKCS. The aim of this study was to investigate morphometric measurements in relation with echocardiographic features of MMVD in American College of Veterinary Internal Medicine (ACVIM) class B1 CKCS [14]. This class, in fact, includes the majority of the breeding population and is very heterogeneous from clinical, morphological and echocardiographic points of view [15]. The identification of phenotypic characteristics associated with more severe forms of MMVD could be useful for setting breeding selection programs aimed at reducing the prevalence of the disease in this breed.

## Material and Methods

### Study population, research question and statistical framework

In this prospective clinical cross-sectional study, we carefully described the morphometry of a small Italian study population of CKCS and then we evaluated the influence of body, thorax, and head dimension on different clinical (heart murmur intensity) and echocardiographic measures/indexes of the severity of MMVD (MVP, regurgitant jet size, and indexed mitral valve and annulus measurements). Furthermore, we investigated the severity of MMVD including a score, assigned according to the degree of MVP, mitral regurgitation jet size and age [8]. To investigate the association between morphometric measures and severity of MMVD we used a method adopted from the causal inference framework [16]. The framework proposes methods to address causal questions accounting for the confounding which affects the association between the exposure and the outcome of interest. In this study we used inverse probability weighting (IPW) analyses that, via weighted regression modelling, adjust for confounders [17-18]. The confounders are used to estimate the probability of being exposed conditional on the values of the confounders and a function of this probability is used to construct weights. The weights are assigned to the subjects in the study population to balance them with respect to the confounders used in the analysis. Balancing subjects in the study population allows to estimate an association that is unbiased from the confounders considered in the analysis while constructing the weights.

### Inclusion Criteria and Clinical Examination

Fifty-two privately owned CKCS with asymptomatic MMVD and no cardiac enlargement (ACVIM stage B1) [14], belonging to different lineages and breeders, were recruited for enrollment in this prospective cross-sectional study. The dogs were examined during breed health screening at the Cardiology Unit of the Veterinary Teaching Hospital - Department of Veterinary Medicine - University of Milan, between April 2019 and June 2020. Informed consent was signed by the owners, according to the ethical committee statement of the University of Milan number 2/2016. All the dogs underwent physical examination, echocardiography, and morphometric evaluation. The data regarding the dates of birth and the genealogy were verified by checking each animal microchip number and family tree in the Italian regional registry and ENCI’s (Ente Nazionale della Cinofilia Italiana) pedigree database. Cardiac auscultation was performed by two well-trained operators (MB and PGB) and the dogs were restrained in standing position in a quiet room by the owners. The detection of a left apical systolic murmur was not considered a mandatory inclusion criterion. The evaluated auscultatory findings were presence/absence, timing, and intensity (0=absent; 1=I-II/VI left apical systolic or soft murmur; 2=III-IV/VI bilateral systolic or moderate and loud murmur respectively; 3=V-VI/VI bilateral systolic or palpable murmur) of murmur [19]. Unless otherwise stated, hereinafter the term murmur refers to a left apical systolic murmur.

The diagnosis of MMVD was based on the echocardiographic evidence of changes of the mitral valve leaflets (thickening and prolapse) and the presence of mitral regurgitation on color-flow Doppler [9]. To be included in the study, dogs must have no evidence of left atrial and left ventricle dilatation, defined as a left atrial-to-aortic root ratio (LA/Ao) ≥ 1.6 on a 2-dimensional echocardiography, and as left ventricular normalized dimensions in diastole (LVIDad) ≥ 1.7, respectively [14]. Blood pressure was indirectly measured with a Doppler method according to the ACVIM consensus statement [20,21].

### Echocardiography and Assessment of Leaflet Measurements, MVP Severity and Jet Size

All echocardiograms were performed by the same operators using a MyLab50 Gold cardiovascular echocardiograph (Esaote, Genova, Italy) equipped with multi-frequency phased array probes (3.5-5 and 7.5-10 MHz), chosen according to the weight of the subject, and with standardized settings. Video clips were acquired and stored using the echo machine software for off-line measurements. The exam was performed according to a standard procedure with concurrent continuous electrocardiographic monitoring [22].

The mitral valve was evaluated using both right and left parasternal long axis 4-chamber views [22,23]. Valve morphology and structures, including the presence/absence and the grade of valvular prolapse, were defined. The right parasternal 4-chamber view was used for the morphological evaluation of the mitral anterior leaflet during its maximum distension in diastole [23]. The anterior mitral valve length (AMVL), width (AMVW), and area (AMVA) were measured, as well as the mitral valve annulus in diastole (MVAd) and systole (MVAs) in the first frame respectively after closing and before opening of the leaflets [23]. All measurements were indexed according to Wesselowski method [23].

Mitral valve prolapse was considered mild if the leaflets were prolapsing but did not cross the line joining their pivotal points (P line), moderate if protruded between the P line and the line joining half of the echoic areas located in the lower part of the atrial septum at the level of atrioventricular junction (T line), severe if the leaflets exceeded the T line [24]. The sphericity index (SI) was calculated as the ratio of the LV long-axis diameter to short-axis diameter in end-diastole from the right parasternal 4-chamber long axis view and a value of SI <1.65 accounted for an increased sphericity and was considered abnormal according to the European Society of Veterinary Cardiology (ESVC) guidelines [25,26], despite the literature did not report SI reference values for small breed dogs, and in particular for CKCS.

The following measurements were taken from the right parasternal short-axis view: LA/Ao obtained in two-dimensional view as described by Hansson et al. [27], and left ventricular diameter measured in M-mode from the short axis with the leading edge to inner edge method at the level of the papillary muscles. Left ventricular normalized dimensions were calculated as described by Cornell et al. (2004) [28]. Left ventricular end-diastolic (EDV) and end-systolic volumes (ESV) were calculated by the Teichholtz method and the values were successively indexed for body surface area (BSA) in order to obtain the end-diastolic (EDVI) and the end-systolic (ESVI) volume indexes [29]. The area length method was used for the calculation of 2-D derived parameters: ejection fraction (EF%), indexed for BSA 2D-EDVI and 2D-ESVI for each patient [29].

The color flow mapping of the mitral valve area was obtained from the left parasternal long axis 4-chamber view [30,31]. A pulse repetition frequency of 5 kHz was used, and the flow gain was adjusted to the maximal level without getting background noise.

The degree of MR (jet size) was assessed using color Doppler and calculating the maximal ratio of the regurgitant jet area signal to left atrium area (ARJ/LAA ratio) [32]. Regurgitant jet size was estimated as the percentage of the left atrial area (to the nearest 5%) that was occupied by the larger jet and it was considered as trivial (<10%, not visible in all the systolic events), trace (<10%, present in all the systolic events), mild (between 10 and 30%), moderate (between 30 and 70%) or severe (>70%) [32,33].

Echocardiographic measurements were performed by one operator (MB) to reduce potential biases. All measurements of interest were repeated on 3 consecutive cardiac cycles, and the mean value was used in the statistical analysis [32]. Within-day intra-observer variability in the studied variables was determined by reanalyzing the parameters measured by the same observer (MB) 3 times after the first measurement on a subset of 10 randomly selected blind exams from the database. The same frames from the same videos were chosen for the evaluation of intra-observer variability. The intra-observer coefficients of variation (CV range in %) were <10% for each tested variable.

For descriptive purpose only, a severity score was assigned according to degree of MVP, mitral regurgitant jet size and age of each subject as reported by Stern et al. in 2015, according to [(Mitral valve prolapse + Regurgitant jet size) x 5] / age formula [34]. Because of the age-related nature of disease severity, a continuous variable was constructed, so that MMVD affecting younger dogs would be considered a more severe disease variant than the one affecting older dogs with the same level of degenerative change. The age of 5 years was chosen as a pivotal point in the breed, where dogs less than 5 years demonstrating clear evidence of disease were considered the most severely affected animals [34,35].

### Morphometrics

The clinical and echocardiographic examination were completed by a specific morphometric evaluation that included the assessment of ENCI’s standard coat color type (Blenheim, ruby, tricolor and black and tan) and the measurement of body, thorax, and head of each dog. All morphometric measurements that were performed and their references points are outlined in Table 1 [36-38]. Body size, cephalic, thoracic and volume indexes were also calculated (Table 1) [36]. During the morphometric evaluation, the dogs were kept calm and in standing position by owners, with the four limbs perpendicular, hand stacking as if they were in exposition. Morphometric evaluation was always performed by the same operator (MB), to reduce systematic errors, on the left side of each dog, to reduce potential biases. Intra-observer variability was < 10%. The circumference of the thorax was measured using a measuring tape. Body and thorax evaluations were performed using a custom-made sliding gauge (Fig 1).

**Table 1.** Morphometric measurements, body indexes and their reference points [35,37-39].

| Body measurements | Thorax measurements | Head measurements |
| --- | --- | --- |
| Height at the withers (WH): distance of the withers from the ground, measured at the top of the shoulder blades | Height (TH): distance between the back and the sternum, measured behind the shoulders | Head length (HL): measured from the top of the occipital ridge to the horizontal line joining the two inner corners of the eyelids |
| Body length (BL): distance between the tip of the shoulder and the tip of the buttock | Width (TW): measured just behind the shoulders | Nose length (NL): measured from the horizontal line joining the two inner corners of the eyelids to the cranial extremity of the truffle |
| Width at chest (CW): measured at the shoulder-humeral joints | Length (TL): distance between the shoulder tip and the midline of the last rib | Head + nose length (HNL): HL + NL |
|  | Circumference (TC<sub>1</sub>, TC<sub>2</sub>, TC<sub>3</sub>)<br>- TC<sub>1</sub> (Anterior or axillary circumference): measured at the level of the anterior part of the axillary cable<br>- TC<sub>2</sub> (Mean or papillary circumference): measured at the level of the first mammary nipples<br>- TC<sub>3</sub> (Lower or basal circumference): measured at the level of the xiphoid process of the sternum, which corresponds above the spinous process of the seventh cervical vertebra | Head width (HW): measured at the zygomatic arches |
|  |  | Head stop angle (HA): angle obtained, with the head seen in profile, by intersection of a line tangent to the frontal region (between the two orbits) and the line of the upper part of the nasal barrel |
| Body indexes |  |  |
| Cephalic index: (HWx100)/HNL |  |  |
| Craniofacial ratio (CFR): NL/HL |  |  |
| Thoracic index: (TWx100)/TH; Height thorax index: (THx100)/WH |  |  |
| Volume index: (Body weightx100)/WH |  |  |
| Body size: (WHx100)/BL |  |  |

**Fig 1.**
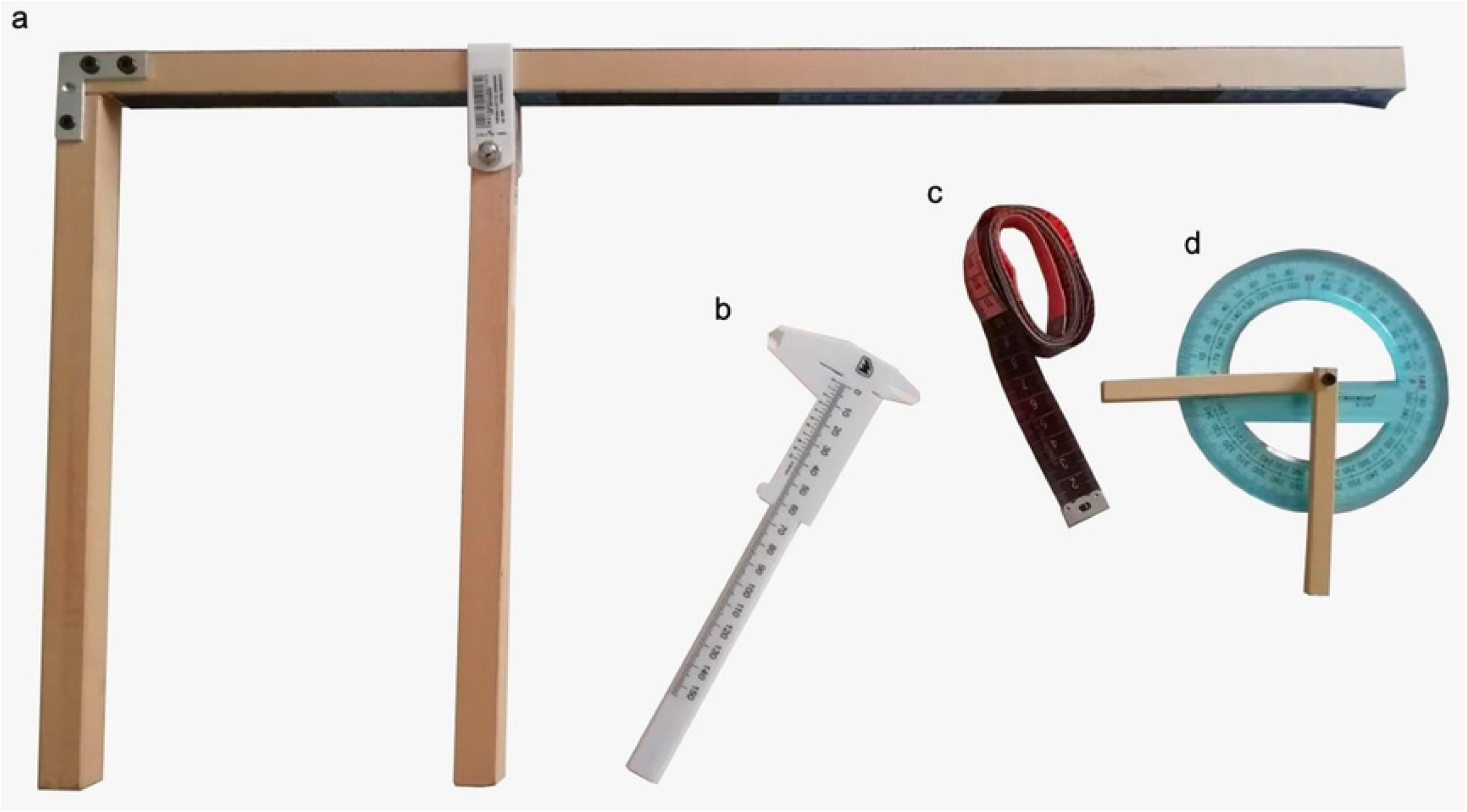
Measurements tools used to evaluate the thorax and the body dimensions. a) The basal part of the custom-made sliding gauge used to measure the height at the withers and width and height of the thorax and the mobile part that, together with the basal part, was used to measure the height at the withers and width and height of the thorax b) The gauge used to measure head length, nose, and head width c) The measuring tape used to measure the three thorax circumferences d) The goniometer used to measure the head’s stop angle.

Supplementary material 1 shows how morphometric measurements were performed and the reference points for each measurement.

The dogs were classified according with the standard values reported by ENCI (https://www.enci.it/media/2405/136.pdf). Standard nose length (NL) was “about 3.8 cm”. It is very interesting to underline that the Italian breed guidelines do not give precise indications regarding the determination of the length of the nose (https://www.enci.it/media/2405/136.pdf). The body condition score (BCS) was recorded for each dog using a 1 to 9 score, and scores 4 and 5 were considered as normal [39].

### Exclusion Criteria

Healthy (n.16) and MMVD affected CKCS at ACVIM stages B2 (n.15), C (n.15) and D (n.13) were not included, as well as those in stage B1 (n.5) with either left atrial or left ventricular enlargement but not both. Dogs with cardiac diseases other than MMVD, such as myocarditis (n.1), congenital heart defects (n.2), cardiac tumors (n.1) and diagnosed arrhythmias (such as supraventricular/atrial premature contractions) (n.4) were not included in the study, as well as subjects with hypertension (n.3) or metabolic diseases (n.5) [21]. Dogs with facial, thoracic and limb malformations (n.2) and dogs younger than one year of age (n.2) were also excluded.

### Statistical Methods

The influence of morphometric measurements on MVP severity, jet size (ARJ/LAA ratio), murmur intensity and echocardiographic indexed measurements (AMVL, AMVW, AMVA, MVAd, MVAs, SI) in the 52 CKCS was evaluated via IPW analyses. An IPW analysis requires to construct two regression models, one for the exposure and one for the outcome. The regression model for the exposure is the propensity score (PS) model which estimates, for all included subjects, the probabilities of being exposed to the risk factor of interest. For this study, the PS model included the 13 morphometric variables (exposures) as a multivariate response (“generalized multivariate propensity scores”) [18], while age, sex, weight, and coat (covariates or confounders) were used as regressors. The probabilities associated to predicted values that resulted from the estimated PS model were the propensity scores estimates. Inverse probability weights were derived as “stabilized inverse probability weights” (SIPW), i.e. the ratio between the multivariate marginal probability density of the exposures and the multivariate probability density of the exposures conditional to covariates [40].

The second regression model is the outcome model, the main analysis model, which aims to estimate the association of the exposures with the response variables. For each type of response variable (6 numerical and 3 ordinal outcomes), normal linear model or ordinal logistic regression models were estimated, with the morphometric variables as exposures, and weighting the observations with the SIPW previously estimated [41]. For the PS model, two alternative models were considered: an additive model (with no interaction between regressors) or an interaction model (with bivariate interactions between some of the regressors). For the PS model and for each of the responses, the bivariate interactions between regressors were explored using normal or ordinal regression models, each of them containing one regressor at the time and only one of the possible bivariate interactions between regressors. Finally, the interaction PS model was estimated adding all bivariate interactions detected as significant in the previous exploratory models. Interaction and additive models were then compared for the best performance. The best PS model between the additive and the one including interactions, was chosen as the model that provided a better balance of the values of the covariates across subjects and a better match of the positivity assumption [16]. The assumption of “positivity” implies that all participants have the potential to receive a particular level of exposure given any value of the confounders. This was checked by identifying the range of values of the exposures where positivity is satisfied as the “multidimensional convex hull” (the smallest convex shape enclosing a given shape) calculated for the observed exposures values [18]. In the outcome model robust standard errors were estimated to take the IPW into account [16].

Outcome models were checked for normality, linearity, variance homogeneity and multicollinearity. Moreover, outcome ordinal models were also checked for violation of the proportional odds assumption, using surrogate residuals [42,43]. The effects of the morphometric indices, that are not linear functions of some single morphometric measurements, were derived from the parameters of outcome models, by means of the “delta method” [44].

All analyses were performed in “R” environment [45].

## Results

### Clinical and Echocardiographic Results

The median age of the included dogs was 4.16 years (IQR_25-75_ 2.91-6): 14 (27%) were younger than 3 years, 25 (48%) between 3 and 6 years, and 13 (25%) older than 6 years. Eleven dogs (21.2%) were intact males, 2 (3.8%) neutered males, 34 (65.4%) intact females, and 5 (9.6%) neutered females. Median weight was 9.15 Kg (IQR_25-75_ 7.80-10.23). Thirty-six subjects (69.2%) weighted more than the proposed breed standard (5-8 Kg), 8 (22.2%) of which were overweight (BCS > 5) and 4 (11.1%) were underweight (BCS < 4). No subjects weighted less than the standard (5 Kg). Neutered females showed higher body weight compared to intact females and intact males (P < .05).

In 26 dogs (50%), no murmurs were found. Soft murmurs (1) were present in 20 dogs (38.5%), whereas in 6 dogs (11.5%) murmurs were of moderate/loud intensity (2). With reference to the 26 dogs with undetectable murmurs, 25 had MVP (19 mild and 6 moderate) and 19 had MR (13 trivial, 3 trace, and 3 mild). Of these 26 subjects, 18 dogs presented MVP and MR, 1 MR only, and 7 MVP only. However, 51 (98.1%) of the 52 included subjects had MVP.

Sphericity index was lower than 1.65 in 48 (92.3%) subjects. Table 2a shows all clinical data (age, body weight and sex), indexed mitral valve measurements and MVP, jet size, murmur severity, and severity score of all included subjects. Moreover, MVAd was larger in subjects older than 6 years than in dogs younger than 3 years (P = .03), whereas MVAs was larger in subjects older than 6 years than in those between 3 and 6 years (P < .001). Sphericity index was lower in subjects older than 6 years compared to subjects with age between 3 and 6 years (P = .01).

**Table 2.**
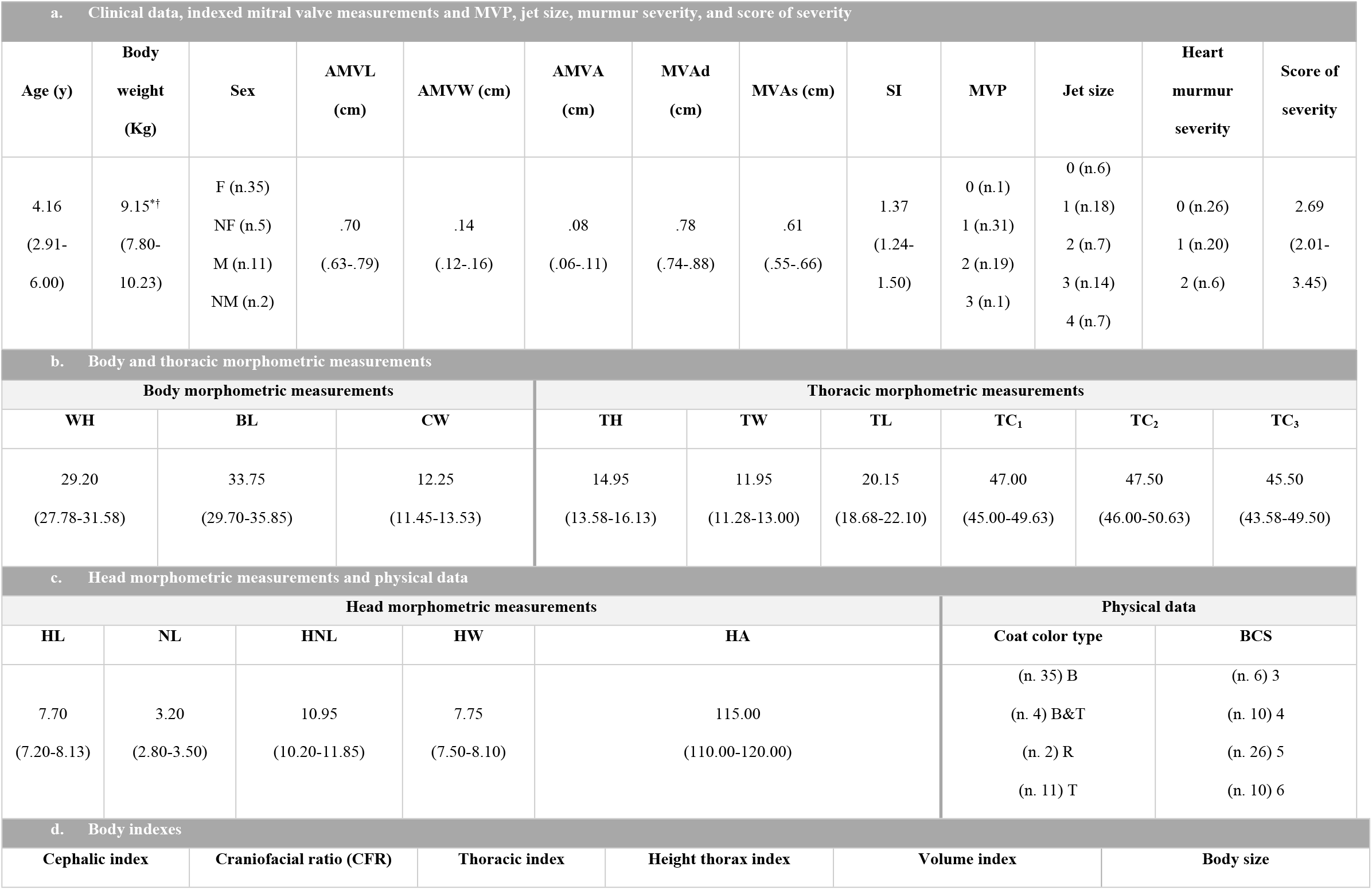
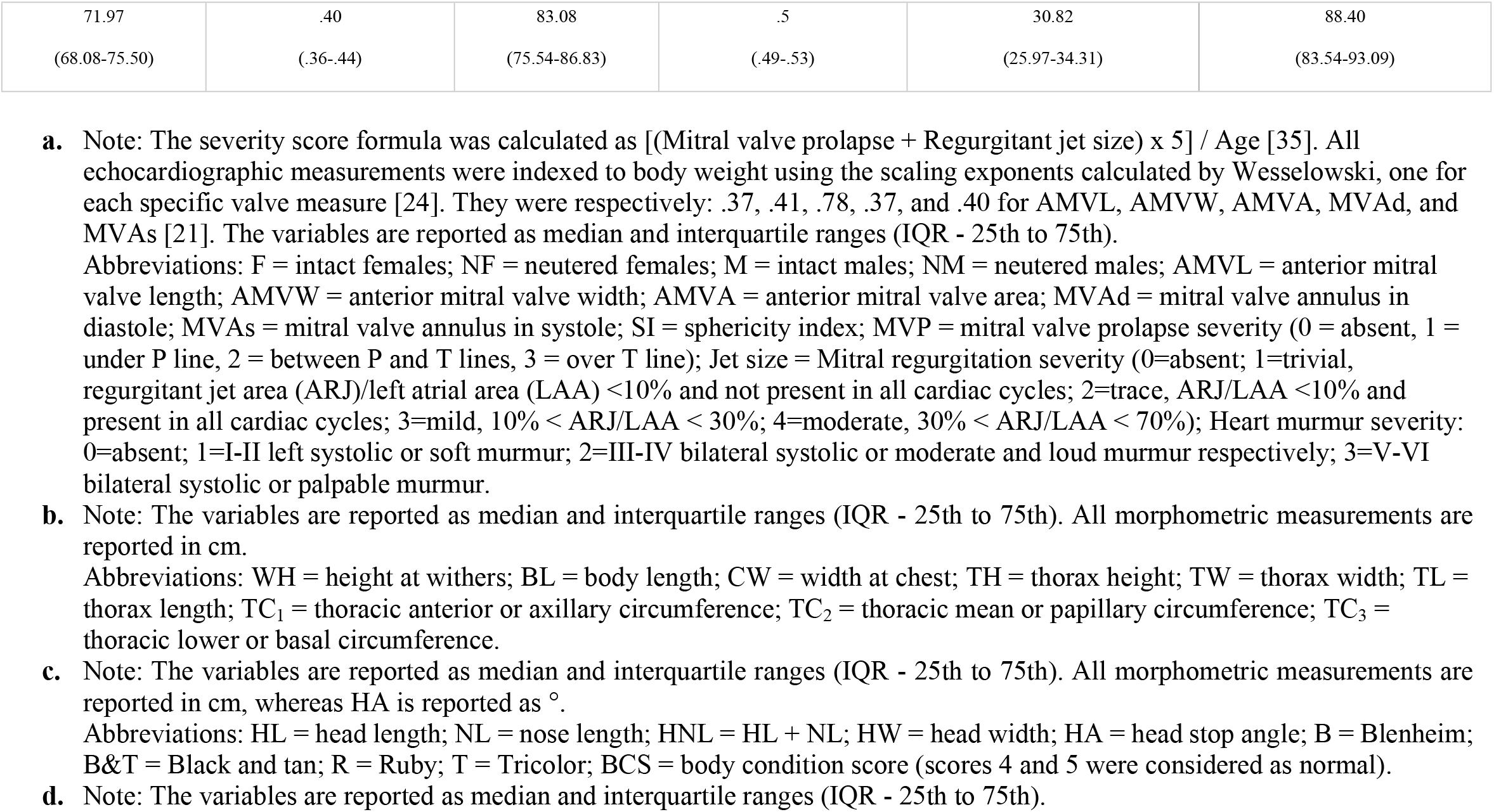
Clinical data, indexed mitral valve measurements, MVP, jet size, murmur severity, severity score and morphometric measurements, indexes, coat color type and BCS of all included subjects.

| a. Clinical data, indexed mitral valve measurements and MVP, jet size, murmur severity, and score of severity |  |  |  |  |  |  |  |  |  |  |  |  |  |  |  |  |
| --- | --- | --- | --- | --- | --- | --- | --- | --- | --- | --- | --- | --- | --- | --- | --- | --- |
| Age (y) | Body weight (Kg) | Sex | AMVL (cm) | AMVW (cm) | AMVA (cm) | MVAd (cm) | MVAs (cm) | SI | MVP | Jet size | Heart murmur severity | Score of severity |  |  |  |  |
| 4.16<br>(2.91-6.00) | 9.15**<br>(7.80-10.23) | F (n.35)<br>NF (n.5)<br>M (n.11)<br>NM (n.2) | .70<br>(.63-.79) | .14<br>(.12-.16) | .08<br>(.06-.11) | .78<br>(.74-.88) | .61<br>(.55-.66) | 1.37<br>(1.24-1.50) | 0 (n.1)<br>1 (n.31)<br>2 (n.19)<br>3 (n.1) | 0 (n.6)<br>1 (n.18)<br>2 (n.7)<br>3 (n.14)<br>4 (n.7) | 0 (n.26)<br>1 (n.20)<br>2 (n.6) | 2.69<br>(2.01-3.45) |  |  |  |  |
| b. Body and thoracic morphometric measurements |  |  |  |  |  |  |  |  |  |  |  |  |  |  |  |  |
| Body morphometric measurements |  |  |  |  | Thoracic morphometric measurements |  |  |  |  |  |  |  |  |  |  |  |
| WH |  | BL |  | CW |  | TH |  | TW |  | TL |  | TC<sub>1</sub> |  | TC<sub>2</sub> |  | TC<sub>3</sub> |
| 29.20<br>(27.78-31.58) |  | 33.75<br>(29.70-35.85) |  | 12.25<br>(11.45-13.53) |  | 14.95<br>(13.58-16.13) |  | 11.95<br>(11.28-13.00) |  | 20.15<br>(18.68-22.10) |  | 47.00<br>(45.00-49.63) |  | 47.50<br>(46.00-50.63) |  | 45.50<br>(43.58-49.50) |
| c. Head morphometric measurements and physical data |  |  |  |  |  |  |  |  |  |  |  |  |  |  |  |  |
| Head morphometric measurements |  |  |  |  |  |  |  |  |  |  | Physical data |  |  |  |  |  |
| HL |  | NL |  | HNL |  | HW |  | HA |  |  | Coat color type |  |  |  | BCS |  |
| 7.70<br>(7.20-8.13) |  | 3.20<br>(2.80-3.50) |  | 10.95<br>(10.20-11.85) |  | 7.75<br>(7.50-8.10) |  | 115.00<br>(110.00-120.00) |  |  | (n. 35) B<br>(n. 4) B&T<br>(n. 2) R<br>(n. 11) T |  |  |  | (n. 6) 3<br>(n. 10) 4<br>(n. 26) 5<br>(n. 10) 6 |  |
| d. Body indexes |  |  |  |  |  |  |  |  |  |  |  |  |  |  |  |  |
| Cephalic index |  |  | Craniofacial ratio (CFR) |  |  | Thoracic index |  |  | Height thorax index |  |  | Volume index |  |  | Body size |  |

|  |  |  |  |  |  |
| --- | --- | --- | --- | --- | --- |
| 71.97<br>(68.08-75.50) | .40<br>(.36-.44) | 83.08<br>(75.54-86.83) | .5<br>(.49-.53) | 30.82<br>(25.97-34.31) | 88.40<br>(83.54-93.09) |

### Morphometric Measurements

Forty-four (84.6%) dogs, 11 (21.1%) males and 33 (63.5%) females, had a height at the withers lower than breed standard (34-36 cm for males and 32-35 cm for females). In 6 (11.5%) subjects the nose length was longer than the standard (3.8 cm), whereas in 45 (86.5%) was shorter than 3.8 cm. In 36 CKCS (69.2%) nose length was shorter than 3.5 cm and in 17 (32.7%) shorter than 3 cm. In only one subject the nose length was equal to the standard measure (3.8 cm). The morphometric measurements, coat color type, and BCS of overall included population are showed in Table 2b and 2c. The morphometric indexes are reported in Table 2d. Neutered females showed greater thorax height, thorax width, TC_3_, and volume index than intact females and greater TC_3_ and volume index compared with intact males (P < .05). Furthermore, intact males showed greater height at withers, head length, and head-nose length compared with intact females (P < .05). No differences between nose length, head width, and head stop angle between sexes (P > .05) were found. Head stop angle was lower (i.e., closer to 90°) in subjects with a weight within standard (5-8 Kg) (P = .04). Subjects with tricolor coat type had larger head width than Blenheim ones (P = .01). Furthermore, younger subjects (<3 years) weighted less than older subjects (P < .001), showed a lower thorax length (P < .05) and volume index (P < .01), as well as higher severity score (P < .05).

### Settings for IPW analysis

The IPW analysis was performed including 49 of the 52 subjects, due to lack of some values, both for ordinal and continuous variables. Furthermore, the covariate “coat color type” has been simplified in “Blenheim” and “other colors” due to the small number of subjects with coat different from Blenheim. For the same reason, the information “neutered/sterilized”, originally incorporated in the covariate “sex”, has not been used.

The PS model with interactions between covariates/confounders provided a better balance in terms of observed confounders between exposure groups (increasing comparability) and accordingly this model was used to build SIPW.

### IPW analyses for ordinal variables

Obtained results were significant for two of the three considered ordinal variables, particularly for heart murmur intensity and jet size. The IPW analysis, in fact, showed that body length (P = .03) and nose length (P < .01) had negative influence on heart murmur intensity (shorter body length and shorter nose were associated to higher murmur intensity). Furthermore, head length (P < .001) had negative influence on jet size (shorter head was associated to larger jet size). However, morphometric measurements had no effects on MVP severity.

The results of the regression analysis for ordinal variables, applied to the included population, are summarized in Table 3a.

**Table 3.** Results of the regression analysis applied to determine the influence of morphometric variables on clinical and echocardiographic parameters in all included subjects.

| Regression analysis |  |  |  |  |  |  |  |  |  |  |
| --- | --- | --- | --- | --- | --- | --- | --- | --- | --- | --- |
|  | BL | TH | TW | TL | TC<sub>1</sub> | TC<sub>2</sub> | TC<sub>3</sub> | HL | NL | HA |
| **a. Ordinal variables** |  |  |  |  |  |  |  |  |  |  |
| **MVP** |  |  |  |  |  |  |  |  |  |  |
| **Jet size** |  |  |  |  |  |  |  | **n |  |  |
| **Heart murmur intensity** | *n |  |  |  |  |  |  |  | **n |  |
| **b. Continuous variables** |  |  |  |  |  |  |  |  |  |  |
| **AMVL** |  |  |  |  |  |  |  | *p |  |  |
| **AMVW** |  |  | *p | *n |  |  |  |  |  |  |
| **AMVA** |  |  |  |  |  |  |  |  |  |  |
| **MVAd** | *p | *n | ***p |  | ***n | ***p | *n | ***p | ***n | ***p |
| **MVAs** |  | **n | *p |  |  |  |  |  |  | ***p |
| **SI** |  |  |  |  | **p |  |  | **p |  |  |

Note: All morphometric measurements are expressed in cm, whereas HA as °. Morphometric variables not influencing clinical and echocardiographic parameters are not reported in the table.
Level of statistical significance: \*\*\* = $P < .001$ ; \*\* = $P < .01$ ; \* = $P < .05$ .
p = positive association; n = negative association.
Abbreviations: MVP = mitral valve prolapse severity; Jet size = severity of mitral regurgitation; Anterior mitral valve: length (AMVL), width (AMVW), area (AMVA); MVAd = mitral valve annulus in diastole; MVAs = mitral valve annulus in systole; SI = sphericity index; BL = body length; TH = thorax height; TW = thorax width; TL = thorax length; TC<sub>1</sub> = thoracic anterior or axillary circumference; TC<sub>2</sub> = mean or papillary circumference; TC<sub>3</sub> = thoracic lower or basal circumference; HL = head length; NL = nose length; HA = head stop angle.

### IPW analyses for continuous variables

Head length (P < .001) had positive influence on anterior mitral valve length (longer head was associated to longer anterior mitral valve leaflet).

Thorax width (P = .01) had positive influence on anterior mitral valve width, whereas thorax length (P = .04) had a negative one: subjects with larger or shorter thorax had thicker anterior mitral valve leaflet.

Body length (P = .02), thorax width (P = .000), mean or papillary circumference (TC_2_) (P < .001), head length (P = .000), and head stop angle (P = .000) had positive influence on mitral valve annulus in diastole, whereas thorax height (P = .02), TC_1_ (P = .000), TC_3_ (P = .02), and nose length (P < .001) had a negative influence.

Thorax width (P = .01) and head stop angle (P = .000) had positive influence on mitral valve annulus in systole, whereas thorax height (P = .002) had a negative influence.

Anterior or axillary thoracic circumference (P = .01) and head length (P = .000) had a positive influence on sphericity index.

The derivation of results of IPW analysis respect to the morphometric indexes showed that only thoracic index was negatively associated with mitral valve annulus in systole and diastole.

The results of the regression analysis for continuous variables, applied to the included population, are summarized in Table 3b.

The performance of the outcome models regarding normality, linearity and variance homogeneity were acceptable according to the diagnostic plots and there were no clues of violation of proportional odds assumption for ordinal models, whereas some multicollinearity problems were detected in the models including interaction terms (data not shown).

## Discussion

Very little is known regarding the relationship between echocardiographic indicators of the severity of MMVD (i.e., MVP severity, jet size and indexed echocardiographic measurements) and the morphometric measures, in all breeds. To the best of our knowledge, this study is the first-ever been done on the relation between morphometric data and echocardiocolor Doppler measures in CKCS. The highlight of any association between morphometric data, severity of echocardiographic lesions, clinical symptoms, and evolution time of the disease will be reached only with a long follow-up of the subjects and with a longitudinal study. With our results we have tried to lay the basis for this scope.

In the present study, is clear that subjects with larger thorax width or shorter thorax length (more barrel-shaped) and shorter head had thicker anterior mitral valve leaflets. Mitral valve annulus in diastole has been observed to be larger in subjects with smaller thorax height (reduced dorso-ventral thoracic dimension), larger thorax width, and greater mean or papillary thoracic circumference (TC_2_). The same was observed in subjects with shorter nose. Regarding mitral valve annulus in systole, the results are superimposable to diastole: annulus has been observed to be greater in subjects with smaller thorax height and larger thorax width. It is also interesting to underline a positive influence of anterior or ancillary thoracic circumference (TC_1_) and head length on sphericity index: the ventricular shape was more spherical in subjects with smaller TC_1_ and shorter head. These findings are similar to that observed for human medicine, in which the MVP is associated with an asthenic habitus, corresponding to a reduced antero-posterior thorax diameter [46,47]. In fact, asthenic habitus in the dog could be considered as a reduction of the ventro-dorsal diameter, given the quadrupedal station. Furthermore, in the study carried out by Olsen in Dachshunds [3], the thoracic circumference was negatively correlated with the severity of MVP. Obviously, from the results obtained, we can only speculate about the influence of some morphometric measures on the heart murmur intensity and the jet size, but not on MVP. However, as observed by Olsen [3], we may suppose that the obtained results may indicate an echocardiographic phenotype more easily associated with mitral valve disease (shorter or thicker anterior mitral valve leaflets, greater mitral valve annulus in systole and diastole, and lower sphericity index).

Only one study in CKCS had shown a negative correlation between the severity of MVP and the body weight, demonstrating that smaller dogs have more severe forms of MMVD [9]. With our results, as stated before, we are not able to demonstrate the same; on the other hand, the association between the cranial morphology of the subjects, the severity of the heart murmur, the jet size dimension, and other valvular characteristics is still relevant. We observed that subjects with shorter head were associated with a higher jet size. Furthermore, subjects with shorter body and nose length had higher heart murmur intensity. Thus, given all the discussed results, CKCS with shorter nose and head and more barrel-shaped thorax would have valvular characteristics related to more severe forms of MMVD than subjects with longer and narrower skull and body. According to our results, the breeding of subjects with cranial morphology tending to brachycephalism (wider and shorter head), may be counterproductive in view of the selected reproduction for MMVD, although additional studies are needed.

In literature, only one study investigated the influence of the coat type (length) on MVP prevalence and severity, particularly in Dachshunds.^3^ No association between MMVD and coat color has ever been described. In the present study, coat color, taken as a single factor, did not affect any of MMVD indexes. Nevertheless, it should be considered that in our sample 68% of the subjects were Blenheim: in Italy this is by far the most common color (65%), as reported by ENCI (https://www.enci.it/libro-genealogico/razze/cavalier-king-charles-spaniel). The association between MMVD and coat color type should be evaluated on a larger population of dogs, including subjects in more advanced ACVIM classes.

Many of the included dogs had mild changes, whose long-term significance is unknown due to the lack of data about follow-up. In this study, as already published, a high percentage of dogs without murmurs had MVP from mild to moderate and we report a high prevalence of echocardiographically detected MR in CKCS with no murmurs, reinforcing the inability of a purely clinical screening to identify the MMVD in this breed.^48^ These proportions evoke the need of additional research to follow-up these subjects and understand which of them will develop the disease with a faster course and which ones will remain stable with mild forms up to advanced age. Furthermore, this will enable to understand if some physical characteristics can be related to the progression of MMVD and, given the results of this study, drive the focus of following research on the evaluation of thorax and skull dimensions of the included subjects.

In the end, the obtained morphometric data, combined with genetic analysis and echocardiographic evaluation of the subjects, could prospectively help to characterize some phenotypes related to more severe forms of MMVD.

Among the strengths of this study, the accurate statistical approach used, based on IPW analyses, allowed to account for the measured confounding variables of the association between morphometric measures and MMVD.

This study had some limitations. The qualitative nature of some echocardiographic parameters should be interpreted with caution. Jet size detected by color flow mapping should only be regarded as a semiquantitative measure of the degree of MR [49]. Several factors such as the quality of the imaging window, the distance to the flow being imaged, gain settings, pulse repetition frequency setting for the color Doppler, the immobility of the patient, and the experience of the operator have an influence on this measure [50]. The left apical 4-chamber view was used because the degree of MR may be underestimated if color flow mapping is performed from the right side of the thorax [49]. Furthermore, it has long been known for humans that, given the 3D morphology of the mitral valve, long axis images that do not include the left ventricular outflow tract (LVOT) greatly overestimate the presence of MVP. Considering recent publications on canine 3D mitral valve morphology [51], both in multi-breed populations and specifically in CKCS, this cannot be ignored any longer. It must be highlighted that many of these dogs may not really have mitral valve prolapse.

To our knowledge, the literature does not report SI cut-off reference values for small breed dogs. Awarded of the limits of the cut-off value chosen, the obtained results suggested that CKCS has a ventricular morphology extremely different from all the breeds described by the guidelines, and so specific SI values are needed for CKCS [25,26].

Eventually, further investigations with a greater number of subjects with coat color type different from Blenheim and in advanced ACVIM stages are needed.

Via IPW analyses, the study attempted to provide an association between morphometric measures and MMVD unbiased from confounding factors (used in the analyses), but the presence of unmeasured relevant confounding is still possible.

## Conclusions

In the CKCS included in the present study, MVP had an epidemiology resembling that known for MVP in humans and Dachshunds [3,46,47]. In fact, MVP severity was significantly positively associated with measures of the degree of MMVD (e.g., jet size, leaflet length, and murmur intensity). In the present study, thorax height had a negative association with AMVL. Furthermore, thorax width and TC_1_ had a positive association on MVAd and SI, respectively.

Surely, the most interesting result obtained is that subjects with shorter head were associated with a higher jet size and subjects with shorter body and nose length had higher heart murmur intensity. Regarding mitral valve and mitral annulus measurements, subjects with more barrel-shaped thorax and shorter nose had shorter and thicker anterior mitral valve leaflets and greater mitral valve annulus in systole and diastole. This means that a brachycephalic morphotype, with dogs much more similar to King Charles Spaniel breed in cephalic morphology, is correlated with a more severe jet size and with valvular characteristics related to worse forms of MMVD and this may be counterproductive in view of the selected reproduction for MMVD. Studying of the B1 subjects’ follow-up and the results of this study would allow for a better understanding of the morphological aspects more often associated with the more severe and/or faster evolution of the disease in the CKCS. This, together with clinical and echocardiographic characterization, could be used as part of a screening program for CKCS defining early selection criteria for the exclusion of a subject from reproduction.

## Acknowledgement

This research received no external funding. The authors are grateful to the many dog owners and breeders for their enthusiastic participation to this work and for their availability.

## S1 Appendix: Demonstration of the instrumental and soft tape measurements

Notes: Body and thorax dimensions were measured using a custom-made sliding gauge. Thoracic circumferences were measured on the dog standing and at rest with a firmly held soft tape measure. Head length, nose length, head and nose length, and head width were measured using a gauge. A goniometer was used to measure the head’s stop angle. The detailed definitions of the measurements are shown in Table 1.33 The dogs here represented were included in the present study and photographed by the authors.

A. WH = height at the withers (cm) (blue line); BL = body length (cm) (yellow line).
B. TH = thorax height (cm) (blue line); TL = thorax length (cm) (yellow line).
C. TC1 = thoracic anterior or axillary circumference (cm) (yellow line); TC2 = thoracic mean or papillary circumference (cm) (green line); TC3 = thoracic lower or basal circumference (cm) (blue line).
D. CW = chest width (cm) (blue line); TW = thorax width (cm) (yellow line).
E. HL = head length (cm) (green line); NL = nose length (cm) (blue line); HNL = head-nose length (cm) (yellow line).
F. HA = head stop angle (°) (green).
G. HW = head width (cm) (red line).

